# Multilevel modelling, prevalence and predictors of hypertension in Ghana: Evidence from Wave 2 of the World Health Organization’s Study on Global AGEing and adult health

**DOI:** 10.1101/751487

**Authors:** Justice Moses K. Aheto, Getachew A. Dagne

## Abstract

**Background:** Hypertension is a major public health issue, a critical risk factor for cardiovascular diseases and stroke, especially in developing countries where the rates remain unacceptably high. In Africa, hypertension is the leading driver of cardiovascular disease and stroke deaths. Identification of critical risk factors of hypertension can help formulate targeted public health programmes and policies aimed at reducing the prevalence and its associated morbidity, disability and mortality. This study attempts to develop multilevel regression, an in-depth statistical model to identify critical risk factors of hypertension.

**Methods:** This study used data on 4381 individuals aged ≥18 years from the nationally representative World Health Organization Study on global AGEing and adult health (SAGE) Ghana Wave 2. Multilevel regression modelling was employed to identify critical risk factors for hypertension based on systolic blood pressure (SBP) (i.e. SBP>140mmHg).

**Results:** The data on 4381 individuals were analysed out of which 27.3% were hypertensive. Critical risk factors for hypertension identified were age, obesity, marital status, health state and difficulty with self-care. Strong unobserved household-level residual variations were found.

**Conclusion:** Hypertension remains high in Ghana. Addressing the problem of obesity, targeting specific interventions to those aged over 50 years, and improvement in the general health of Ghanaians are paramount to reducing the prevalence and its associated morbidity, disability and mortality. Lifestyle modification in the form of dietary intake, knowledge provision supported with strong public health message and political will could be beneficial to the management and prevention of hypertension.

## Background

Hypertension remains one of the biggest threats to public health globally, especially in the low- and middle-income countries where the prevalence is the highest as a result of more people residing in these countries, and with greater priority and interest in infectious diseases [1, 2]. It contributes significantly to the global burden of cardiovascular and its related illnesses like stroke, kidney and heart failures, and their resultant premature morbidity, disability and mortality. Hypertension is responsible for 51% of deaths from stroke and at least 45% from heart diseases and overall, it is responsible for about 50% of deaths from stroke and heart disease [1, 3]. The Global Burden of Disease Study Group reported that in 2017, cardiovascular diseases were the leading cause of death in Africa and were responsible for 1.42 million deaths or 16.4% of the total deaths in all ages [4]. The number of deaths represent 61.0% increase over the estimated number of cardiovascular deaths in 1990. High blood pressure, one of the Non-Communicable Diseases (NCDs), is the leading risk factor for deaths in Africa responsible for nearly two-thirds of the cardiovascular deaths in the region. Globally, Africa has the highest prevalence of high blood pressure (27%) [5] and a common cause of medical hospitalization in the region [6] and responsible for over 50% of first time acute stroke [7, 8]. Previous studies reported that with the ageing population and the rising urbanization and its associated stress, sedentary lifestyle and ‘Western’ diet, high blood pressure will continue to rise [2, 8–10].

Previous studies observe that older age, race, cigarette smoking, high salt intake, high body mass index (BMI), alcohol use, female sex, urban residence, physical inactivity and genetics are among the main factors associated with hypertension [8, 11–16]. Though not a significant problem previously in groups like young and rural populations, hypertension is now a critical public health problem in these groups [8, 17–19].

There is a renewed political will to address NCDs as a result of the third high-level meeting of the United Nations General Assembly in October 2018 during which Heads of State and Governments committed to reorienting health systems to respond to the needs of the rapidly ageing population in relation to NCDs [8, 20, 21]. However, sound statistical methods are required to analyse the barriers and facilitators of hypertension in order to achieve this ambitious goal. It is against this background that this study attempts to estimate the prevalence and to employ a multilevel regression model to identify critical risk factors for hypertension to inform sound and targeted policies aimed at improving cardiovascular health outcomes among this population.

## Methods

### Data source and study population

This work was based on the nationally representative World Health Organization Study on global AGEing and adult health (SAGE) Ghana Wave 2 conducted in the period 2014/2015. SAGE is a multi-country study that collects data to complement existing ageing data sources to inform policy and programmes. Information on subjective wellbeing, risk factors and preventive health behaviours, quality of life and health care utilization, perceived health status, household characteristics, socio-demographic and social cohesion were collected from individuals aged 50 years. A smaller sample of those aged 18-49 years were also included in the study. Detailed description of the methods is published elsewhere [22].

### Outcome variable

The outcome variable of interest in the study is hypertension status based on systolic blood pressure measurement (SBP>140mmHg).

### Covariates

This study considered several covariates based on literature on factors influencing hypertension, including other potential health variables yet to be established in the literature as risk factors. These include age, obesity, ethnicity, sex, marital status, type of toilet facility in household, health state and difficulty with self-care at the time of the interview, alcohol consumption, type of cooking fuel, household wall and floor types [2, 11-14, 23-25].

### Statistical analysis

Descriptive statistics were used to summarize the distribution of selected background characteristics of respondents. Further analyses were conducted to examine individual and household-level factors that might be significantly associated with hypertension and explored unobserved household level effects on the outcome. Both single-level and multilevel (mixed effects) logistic regression models were applied on 4381 individuals residing in 3790 households with complete measurements on hypertension as well as complete measurements on potential explanatory variables considered in the final models. The extension of the single level logistic regression model to the multilevel logistic regression model is warranted because of the hierarchical structure of the SAGE dataset where we have individuals nested within households. Specifically, we applied multilevel logistic regression models to examine possible differences in hypertension among individuals across households while simultaneously identifying potential risk factors. Thus, the multilevel modelling approach [26] placed particular emphasis on household level differences in the risk of hypertension among individuals and the extent of nesting of hypertension within a household which cannot be achieved through a single level logistic regression model.

For quantifying the proportion of total variation attributable to within-households differences, the household-level Variance Partition Coefficient (VPC) [27] were employed, which is defined as VPC = (household-level variance/ (household-level variance + individual-level variance)). The individual-level residual is assumed to follow a standard logistic distribution with mean zero and variance π^2^ /3, where π = 3.14 [28].

Model parameters were obtained using maximum likelihood. Identity covariance structure provided a good fit to the data in the multilevel logistic model. The goodness of fit for the fitted models was examined using a likelihood ratio test (LRT), Akaike Information Criterion (AIC) and Bayesian Information Criterion (BIC). Variance inflation factor (VIF) was used to check multicollinearity, and a VIF value below 10 was considered acceptable [29]. All the analyses were performed using STATA Version 14 [30]. Backward elimination was employed to select candidate set of risk factors for multivariable logistic regression analysis. P-value < 0.05 was used to declare statistical significance.

### Ethical approval and consent to participate

SAGE was approved by the World Health Organization’s Ethical Review Board (reference number RPC149) and the Ethical and Protocol Review Committee, College of Health Sciences, University of Ghana, Accra, Ghana. Written informed consent was obtained from all study participants. All methods were performed in accordance with the relevant guidelines and regulations.

## Results

### Participant characteristics

Out of the 4381 individual respondents, 1273 (27.3%) had hypertension. Approximately 76% of the respondents were at least 50 years old and 58.9% were females. Over 48% of the respondents were Akan, with the Guan ethnicity in the minority (4.1%) and over 55% reported being currently married and about 74% had fathers with no formal education. Over 12% were obsess and over 29% reported ever using a tobacco while over 32% reported that they did not know high salt diet can cause health problems. Household characteristics included durable material for walls (62.8%), non-flush toilets (84.6%), shared toilets (76.9%) and hard floor (85.1%) (Table 1).

**Table 1:**
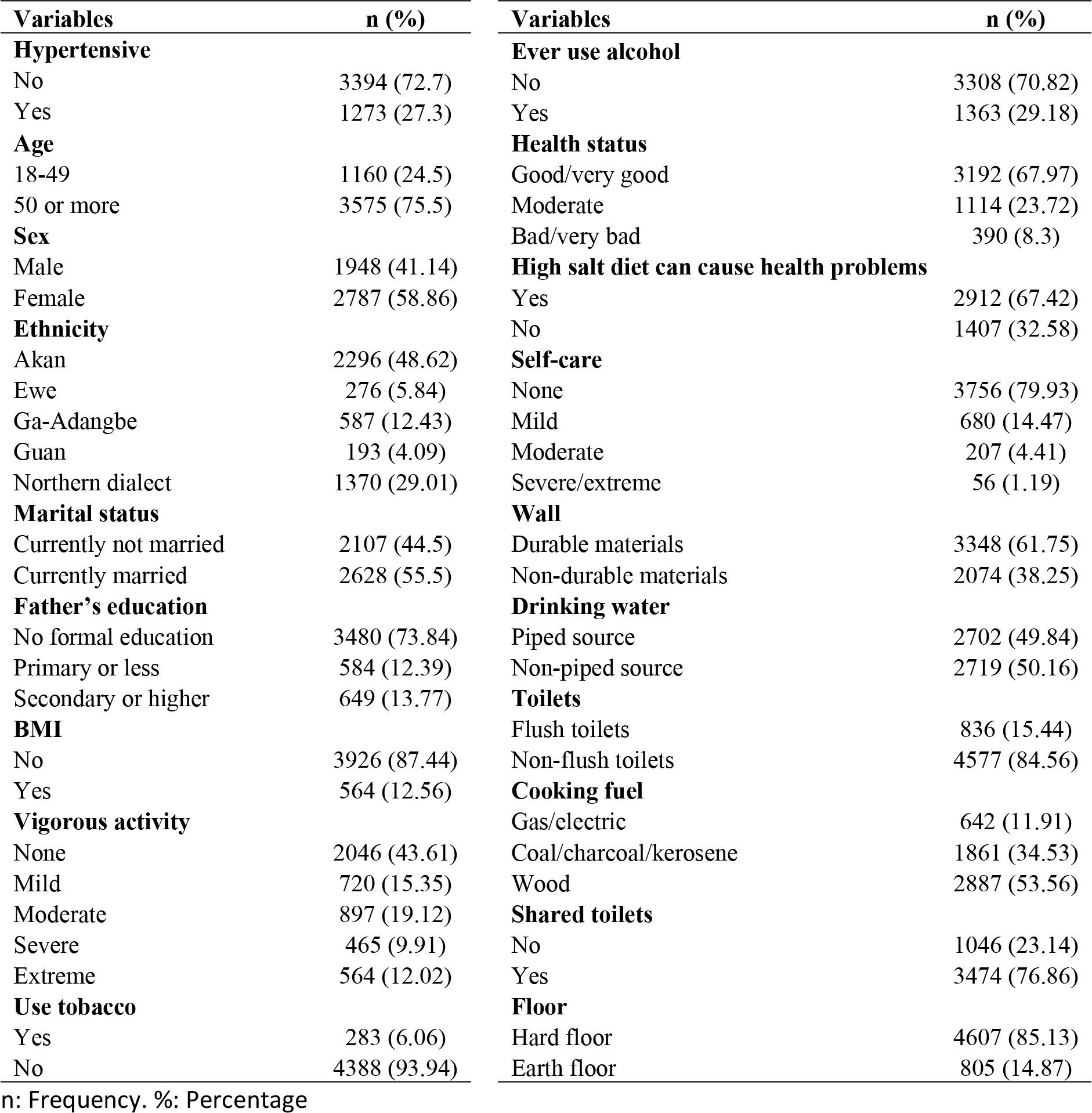
Summary of selected background characteristics of respondents

### Predictors of hypertension

The univariable analyses identified age, obesity, sex, marital status, toilet facility, health state, difficulty with self-care, wall type, floor type and ethnicity as significant predictors of hypertension. Significant predictors of hypertension in the multivariable model include age, obesity, marital status, toilet facility, health state and difficulty with self-care.

Comparing the single-level multivariable logistic regression (Table 2) to the multilevel logistic regression model (Table 3), the multilevel model provided a good fit to the data. Thus, the multilevel logistic regression is preferred to the single-level multivariable model.

**Table 2:** Risk factors for hypertension from single-level logistic regression models

| Variables | Single-level univariable logistic |  | Single-level multivariable logistic |  |
| --- | --- | --- | --- | --- |
|  | UOR (95% CI) | P-value | AOR (95% CI) | P-value |
| **Age** |  | <0.001*** |  |  |
| <50 years | ref |  | ref |  |
| 50 or more | 5.12 (4.13-6.35) | <0.001*** | 4.85 (3.85-6.11) | <0.001*** |
| **Obesity** |  | <0.001*** |  |  |
| No | ref |  | ref |  |
| Yes | 1.51 (1.25-1.83) | <0.001*** | 1.47 (1.19-1.83) | <0.001*** |
| **Sex** |  | <0.001*** |  |  |
| Male | ref |  | ref |  |
| Female | 1.28 (1.12-1.46) | <0.001*** | 1.11 (0.94-1.31) | 0.209 |
| **Marital status** |  | <0.001*** |  |  |
| Not currently married | ref |  | ref |  |
| Currently married | 0.72 (0.63-0.82) | <0.001*** | 0.76 (0.66-0.89) | <0.001*** |
| **Toilet facility** |  | <0.001*** |  |  |
| Flush toilets | ref |  | ref |  |
| Non-flush toilets | 0.74 (0.62-0.88) | 0.001** | 0.79 (0.63-0.99) | 0.039* |
| **Health state** |  | <0.001*** |  |  |
| Good/ very good | ref |  | ref |  |
| Moderate | 1.91 (1.65-2.22) | <0.001*** | 1.34 (1.14-1.58) | <0.001*** |
| Bad/very bad | 2.08 (1.66-2.6) | <0.001*** | 1.32 (1-1.73) | 0.047* |
| **Difficulty with self-care** |  | <0.001*** |  |  |
| Good/ very good | ref |  | ref |  |
| Mild | 1.18 (0.98-1.41) | 0.076 | 1.05 (0.86-1.28) | 0.646 |
| Moderate | 2.13 (1.59-2.85) | <0.001*** | 1.57 (1.09-2.25) | 0.015* |
| Severe/extreme | 3.32 (1.96-5.65) | <0.001*** | 1.78 (0.77-4.13) | 0.177 |
| **Alcohol** |  | 0.605 |  |  |
| No | ref |  | ref |  |
| Yes | 1.04 (0.9-1.2) | 0.605 | 1.09 (0.92-1.28) | 0.317 |
| **Cooking fuel** |  | 0.368 |  |  |
| Gas/ electric | ref |  | ref |  |
| Coal/charcoal/kerosene | 0.96 (0.77-1.18) | 0.681 | 0.97 (0.75-1.26) | 0.816 |
| Wood | 0.88 (0.72-1.09) | 0.24 | 0.87 (0.66-1.13) | 0.296 |
| **Walls** |  | 0.005** | - |  |
| Durable materials | ref |  |  |  |
| Non-durable material | 0.82 (0.72-0.94) | 0.005** |  |  |
| **Floor** |  | 0.026* | - |  |
| Hard floor | ref |  |  |  |
| Earth floor | 0.8 (0.66-0.97) | 0.026* |  |  |
| **Ethnicity** |  | <0.001*** | - |  |
| Akan | ref |  |  |  |
| Ewe | 1.33 (1.01-1.74) | 0.04* |  |  |
| Ga-Adangbe | 1.31 (1.08-1.6) | 0.007** |  |  |
| Guan | 0.8 (0.56-1.13) | 0.203 |  |  |
| Northern dialect | 0.87 (0.75-1.02) | 0.08 |  |  |

UOR: unadjusted odds ratio. AOR: adjusted odds ratio. CI: confidence interval. ref: reference category. \*: P-value <0.05. \*\*: P-value<0.01. \*\*\*: P-value <0.001

**Table 3:** Risk factors for hypertension from multilevel (mixed effect) logistic regression models

| Variables | Mixed effect logistic |  | Parameter (95% CI) |
| --- | --- | --- | --- |
|  | AOR (95% CI) | P-value |  |
| **Age** |  |  |  |
| <50 years | ref |  |  |
| 50 or more | 5.4 (4.11-7.09) | <0.001*** |  |
| **Obesity** |  |  |  |
| No | ref |  |  |
| Yes | 1.51 (1.19-1.91) | 0.001** |  |
| **Sex** |  |  |  |
| Male | ref |  |  |
| Female | 1.11 (0.94-1.33) | 0.225 |  |
| **Marital status** |  |  |  |
| Not currently married | ref |  |  |
| Currently married | 0.75 (0.64-0.89) | 0.001** |  |
| **Toilet facility** |  |  |  |
| Flush toilets | ref |  |  |
| Non-flush toilets | 0.79 (0.62-1.01) | 0.06 |  |
| **Health state** |  |  |  |
| Good/ very good | ref |  |  |
| Moderate | 1.38 (1.15-1.65) | <0.001*** |  |
| Bad/very bad | 1.35 (1-1.83) | 0.047* |  |
| **Difficulty with self-care** |  |  |  |
| None | ref |  |  |
| Mild | 1.06 (0.85-1.32) | 0.62 |  |
| Moderate | 1.64 (1.1-2.44) | 0.016* |  |
| Severe/extreme | 1.88 (0.74-4.73) | 0.182 |  |
| **Alcohol** |  |  |  |
| No | ref |  |  |
| Yes | 1.1 (0.92-1.31) | 0.282 |  |
| **Cooking fuel** |  |  |  |
| Gas/ electric | ref |  |  |
| Coal/charcoal/kerosene | 0.96 (0.73-1.28) | 0.799 |  |
| Wood | 0.84 (0.63-1.14) | 0.266 |  |
| **Individual-level variance** |  |  | 3.29 |
| **Household-level variance** |  | 0.022 | 0.46 (0.15 – 1.48) |

AOR: adjusted odds ratio. CI: confidence interval. ref: reference category. \*: P-value <0.05. \*\*: P-value<0.01. \*\*\*: P-value <0.001

Significant predictors of hypertension in the multilevel model include age, obesity, marital status, health state and difficulty with self-care.

Significant unobserved household-level variations in hypertension were found. The results from the variation analyses showed that over 12% of variance in hypertension could be attributable to residual household-level variations after adjusting for individual and household level factors considered in the multilevel model.

Individuals aged ≥50 years were at increased risk of hypertension compared to those aged 18-49 years (OR=5.4, 95% CI: 4.11-7.09). There is a 51% increase in the odds of hypertension among individuals who are obese compared to their counterparts who are not obese (OR=1.51, 95% CI: 1.19-1.91). Individuals who are currently married had 25% less odds of having hypertension compared to those who are not currently married (OR=0.75, 95% CI: 0.64-0.89). Individuals who rated their health state as moderate or bad/very bad had 38% and 35% higher odds of having hypertension, respectively compared to those who rated themselves as good/very good (OR=1.38, 95% CI: 1.15-1.65 and OR=1.35, 95% CI: 1.0-1.83). Those who rated themselves as having moderate difficulty with self-care had 64% higher odds of having hypertension compared to those who rated themselves as having no difficulty OR=1.64, 95% CI: 1.1-2.44) (Table 3).

## Discussion

The study sets out to estimate hypertension prevalence and to develop a novel multilevel logistic regression model to identify critical risk factors of hypertension to help in formulating targeted policies that could improve cardiovascular health among the Ghanaian adults. In this study, a hypertension prevalence of 27.3% was observed, suggesting that hypertension among Ghanaian adults is still a serious public health issue. This prevalence exceeded the 18% prevalence observed in her neighbouring country Burkina Faso [25], 24.5% in Kenya [31], and 8% in Tanzania [32] but lower than the 31% observed in Nigeria [33]. Critical risk factors independently associated with hypertension while adjusting for the unobserved household-level effects were age, obesity, marital status, health state and difficulty with self-care.

Of critical importance to this study is the quantification of residual household-level effects on hypertension among adults, which represent differences in household-level hypertension outcomes that cannot be explained by the available household-level covariates. Generally, the health and the general wellbeing of individuals is heavily reliant on the households they belong to. Thus, the households determine the resources, opportunities and risks available to the individual over their life course [34-36]. We observed strong residual household-level variations in hypertension and that over 12% of variation in hypertension in adults could be attributable to unobserved household-level variations after adjusting for the risk factors in the model. This could be as a result of household-level, social and environmental factors not considered in our model.

The study broadly supports earlier studies that examined determinants of hypertension in developing countries. For instance, older age group, obesity, rating health state as moderate or bad/very bad, and rating level of difficulty for self-care as moderate were associated with increased odds of hypertension in adults, and those who were currently married had reduced odds of hypertension [3, 11-15, 25, 33, 37, 38]. The association between hypertension and the older age group observed in this study could be attributable to variations in the arterial structure and function, notably arterial stiffening with adverse consequences on cardiac structure and function [33, 39, 40]. Individuals who were currently married had reduced odds of hypertension compared to those who were not currently married. Protective effects for marriage on health has been established in previous studies [41, 42]. However, this finding is not consistent with a previous study that observed that married women had increased odds of hypertension but no such association was found in men in the same study [3]. It also contradicts a previous study [43] that did not find an association between marital status and hypertension but this could be as a result of the setting or how the variable marital status was categorised. Marital status is a critical social characteristic which is well known to predict a range of health outcomes such as cardiovascular illnesses [44, 45] and mortality in general [42, 46].

Obesity, which is one of the modifiable risk factors considered in this study showed an association with hypertension. Individuals who were obese had higher odds of hypertension compared to their counterparts who were not obese, a finding which is consistent with previous studies [25, 31-33, 38, 47]. The association between hypertension and obesity had been established and well known, and reducing BMI is part of the advice provided in the treatment of hypertension [25, 48]. This could be due to metabolic and endocrine disorders as a result of increasing BMI [49].

Individuals who rated their health state as moderate or bad/very bad on the day of the interview had increased odds of hypertension compared to those who rated their health state as good/very good. Also, those who rated their difficulty with self-care as moderate had increased odds of hypertension compared to their counterparts who rated themselves as having no difficulty. These findings are plausible because in a previous study, a hypertensive group had significantly lower age-adjusted health status scores compared to non-hypertensive group [50]. To the best of our knowledge, this is the first study to have established an association between hypertension and health variables like self-reported health state and difficulty with self-care. The significant association between hypertension and these health variables suggest the need to improve the overall health of Ghanaian adults to reduce and prevent the high prevalence of hypertension among this group.

One of the major public health issues in Ghana presently is the rise in the prevalence of NCDs [51, 52]. The findings in the present study provided vital and current information on prevalence and critical risk factors for hypertension that can be used by policymakers and health practitioners for better understanding of hypertension and its prevention and management, which could lead to more effective prevention approaches, patient management and improved cardiovascular outcomes. The study highlights the need to address the problem of obesity, targeting specific interventions to those aged over 50 years, and improvement in the general health of the Ghanaian population as a primary intervention is warranted as part of an overall strategy to reduce the hypertension prevalence and its resultant premature morbidity, disability and mortality.

### Strength and limitation of the study

The strengths of this study include the fact that it utilised data from a nationally representative population-based survey which is globally respected for its sound survey methods and sound data quality on individuals, their households and communities in which they reside. The large samples drawn nationwide permits generalization of findings to the population of adults in Ghana and that of adults from other similar populations globally. The study also used a novel multilevel modelling approach, permitting the study of unobserved household-level effects on hypertension. Thus, providing much more information about why individuals from certain households are more likely to be hypertensive while others are not and at the same time investigating underlying associations between hypertension and the risk factors which could not have been possible using single-level logistic regression approach. Despite these strengths, the study has limitations and so the findings should be interpreted with caution. For example, the analytical techniques employed in the analysis of the data could not establish cause and effect relationship between hypertension and the risk factors considered. Also, some of the risk factors such as health state and difficulty with self-care were based on self-reports and so could introduce reporting bias.

## Conclusion

Findings from the study show that prevalence of hypertension remains high among Ghanaian adults. This study developed a novel multilevel binary logistic regression model which captures unobserved household-level effects and identified critical risk factors of hypertension which can aid formulation of health policies and intervention strategies that will improve cardiovascular health outcomes of the Ghanaian adults. Lifestyle modification in the form of dietary intake, knowledge provision supported with strong public health messages and political will could be beneficial to the management and prevention of hypertension. Active screening for hypertension should be encouraged to identify undiagnosed cases to minimise the danger of stroke and cardiovascular diseases, but there is the need to improve the health systems and services in the country to reap the full benefits of such interventions. There is also the need to target younger populations to minimize their risk of developing hypertension during adulthood/old age. Further study to identify as-yet unidentified risk factors that might account for the substantial unexplained household-level variations in adult hypertension is warranted.

## Abbreviations

AIC: Akaike Information Criterion
BIC: Bayesian Information Criterion
BMI: Body Mass Index
LRT: Likelihood Ratio Test
NDCs: Non-Communicable Diseases
SBP: Systolic Blood Pressure
VIF: Variance Inflation Factor
VPC: Variance Partition Coefficient

## Author contributions

JMKA developed the concept and analysed the data and wrote the first draft manuscript. JMKA and GAD contributed to the writing and reviewing of the various sections of the manuscript. All the authors reviewed the final version of the manuscript before submission. All authors read and approved the final manuscript.

## Acknowledgements

This Fellowship was supported by the University of Ghana Building a New Generation of Academics in Africa (BANGA-Africa) Project with funding from the Carnegie Corporation of New York. The statements made and views are solely the responsibility of the authors. We are also grateful to Prof. Alfred Yawson and his Ghana Team and all respondents and interviewers who made the SAGE survey in Ghana possible. Financial support was provided by the US National Institute on Aging through Interagency Agreements (OGHA 04034785; YA1323-08-CN-0020; Y1-AG-1005-01) with the World Health Organization and a Research Project Grant (R01 AG034479-64401A1). WHO contributed financial and human resources to SAGE. The Ministry of Health, Ghana, is supportive of SAGE. The University of Ghana’s Department of Community Health contributed training facilities, data entry support, and storage of materials. The Ghana Statistical Office provided the sampling information for the sampling frame and updates.

## Consent for publication

Not applicable

## Data availability

Data is freely available upon making official request to WHO-SAGE Team through the WHO website at http://www.who.int/healthinfo/sage/cohorts/en/.

